# Structural basis of RNA polymerase III transcription repression by Maf1

**DOI:** 10.1101/859132

**Authors:** Matthias K. Vorländer, Florence Baudin, Robyn D. Moir, René Wetzel, Wim J. H. Hagen, Ian M. Willis, Christoph W. Müller

## Abstract

Maf1 is a highly conserved central regulator of transcription by RNA polymerase III (Pol III), and Maf1 activity influences a wide range of phenotypes from metabolic efficiency to lifespan. Here, we present a 3.3 Å cryo-EM structure of yeast Maf1 bound to Pol III, which establishes how Maf1 achieves transcription repression. In the Maf1-bound state, Pol III elements that are involved in transcription initiation are sequestered, and the active site is sealed off due to ordering of the mobile C34 winged helix 2 domain. Specifically, the Maf1 binding site overlaps with the binding site of the Pol III transcription factor TFIIIB and DNA in the pre-initiation complex, rationalizing that binding of Maf1 and TFIIIB to Pol III are mutually exclusive. We validate our structure using variants of Maf1 with impaired transcription-inhibition activity. These results reveal the exact mechanism of Pol III inhibition by Maf1, and rationalize previous biochemical data.

## INTRODUCTION

RNA polymerase III transcribes tRNAs, 5S ribosomal RNA, U6 spliceosomal RNA and other small, folded RNAs involved in fundamental cellular processes. Among these products, tRNAs are numerically the most abundant RNAs in the cell and their synthesis, along with 5S rRNA, is energetically costly, consuming an estimated 15% of the nucleotides used by dividing cells for nuclear gene transcription^1^. With its central role in the expression of genes required for cell growth, upregulation of Pol III transcription is increasingly recognized as a prerequisite for oncogenic transformation^2–5^. Pol III transcription is hence tightly regulated in response to stress and growth signals, and several signaling pathways converge to modulate the function of the Pol III machinery including its central negative regulator, Maf^2,6^. Maf1 is conserved across eukaryotes, and has been studied in yeast, flies, worms, plants, parasites and mammalian systems (reviewed in^7,8^). Recently, it was demonstrated that overexpression of Maf1 in the gut of flies increases the lifespan of these animals^9^. Effects of Maf1 on lifespan have also been reported in *C. elegans* although in this case it is the loss of Maf1 function that extends lifespan. Overexpression of the *C. elegans* ortholog MAFR-1 shortens lifespan in *mafr-1* mutant worms but has no effect on the longevity of wild-type worms^10^. Whole body knockout of Maf1 in mice results in a striking lean body composition and resistance to diet-induced obesity^11^. These phenotypes are due in part to metabolic inefficiencies that are thought to be driven by a futile cycle of tRNA synthesis and degradation that wastes cellular energy and reprograms central metabolic pathways^11–13^. The ability of Maf1 to affect body composition and metabolism under this model is supported by its function in mammals as chronic Pol III repressor whose activity can be enhanced by limiting nutrients and stressors^13,14^. Maf1 expression also promotes the differentiation of mouse embryonic stem cells into mesoderm and the differentiation of NIH3T3-L1 cells into adipocytes^15^. These studies demonstrate the central role of Pol III transcription and its inhibition by Maf1 in diverse aspects of organismal physiology.

Maf1 activity is regulated by phosphorylation in response to stress and nutrient availability^1,6^. Phosphosites are located in an internal, poorly conserved region connecting conserved segments at the N- and C-terminus. Dephosphorylated Maf1 accumulates in the nucleus and binds Pol III^1^. Despite its central role in Pol III regulation, the precise mechanism of Maf1-mediated repression remains elusive. Crystal structures have been determined of human^16^ and plant^17^ Maf1, and a low resolution (~20 Å) electron-microscopy reconstruction of Maf1 bound to Pol III was described^16^. Here, we present a 3.3 Å cryo-EM reconstruction of the *Saccharomyces cerevisiae* Maf1-Pol III structure, which differs from the previous proposed model of Maf1 binding and explains the molecular mechanism of Pol III inhibition by Maf1.

## RESULTS

### Cryo-EM structure of the Maf1-Pol III complex

We assembled a complex of purified endogenous *S. cerevisiae* Pol III and a recombinant Maf1 construct lacking an internal non-conserved region that is predicted to be disordered (Maf1-id, Δ64-194). Previous work has shown that the Maf1-id protein is phosphoregulated normally and is active in Pol III repression *in vivo*^18^. Importantly, the recombinant protein shows improved solubility at low salt, which was critical for the formation of a stable Maf1-Pol III complex. Sample crosslinking was performed to prevent complex dissociation during the preparation of cryo-grids, and the complexes were imaged on a Titan Krios microscrope with a K2 detector. A cryo-EM reconstruction at 3.25 Å was determined from 117,442 particles (Figures S1 and S2) and allowed the building of an atomic model (Figures 1 and S1). The Pol III core was resolved at better than 3 Å, while local resolution for Maf1 density varied between 3.6 and 4.3 Å (Figure S1). Yeast Maf1 contains a central beta sheet sandwiched by three helices on one side (interface A) and a single helix and two loops on the other side (interface B) (Figure 2) and is very similar to human Maf1^16^ with a C_α_ rmsd of 2.4 Å, despite the large evolutionary distance between yeast and human.

**Figure 1.**
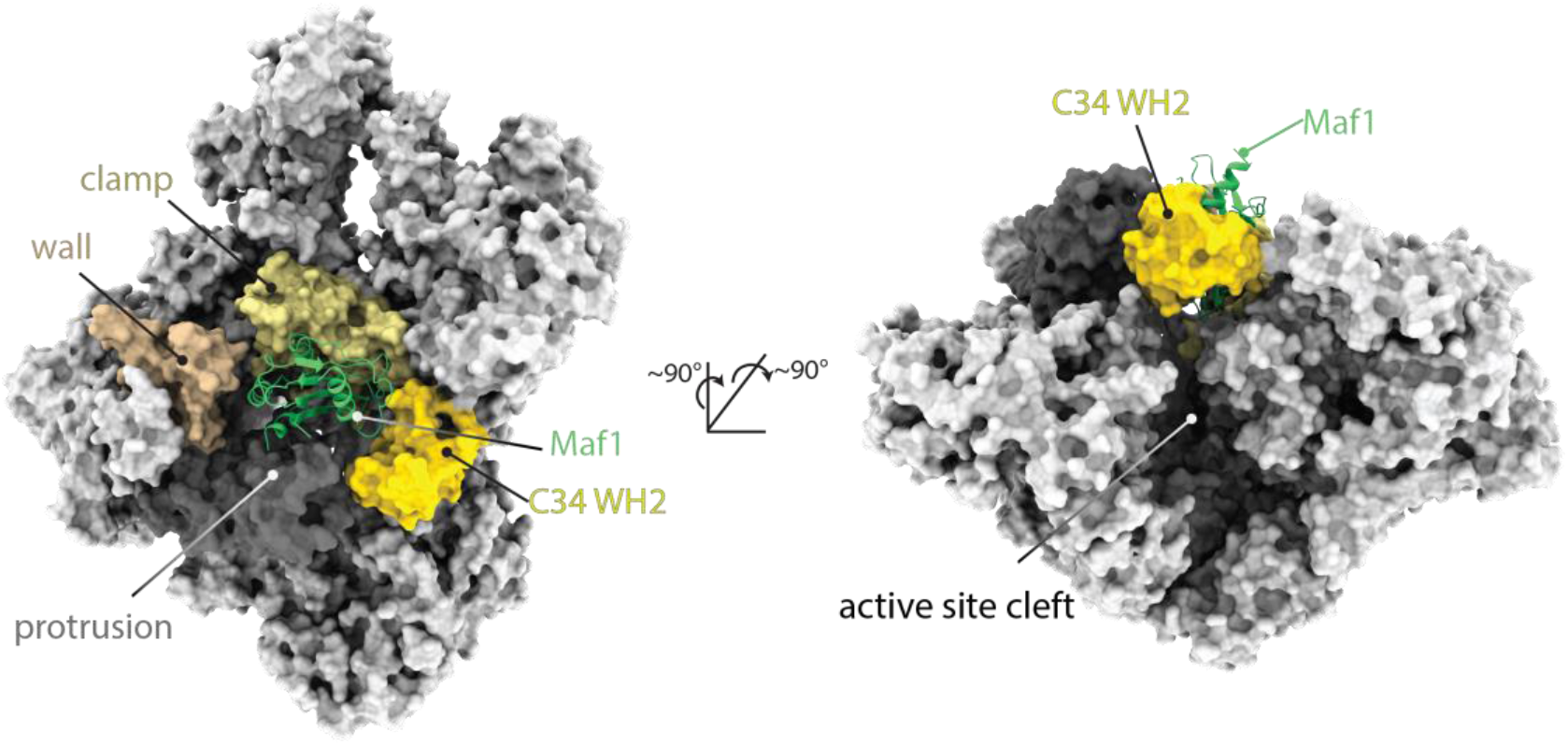
Structure of the Maf1-Pol III complex. Pol III is shown as a surface and Maf1 in ribbon representation. Maf1 (green) is bound between the clamp (ochre), wall (beige), protrusion (dark grey) and C34 WH2 domain (yellow).

**Figure 2.**
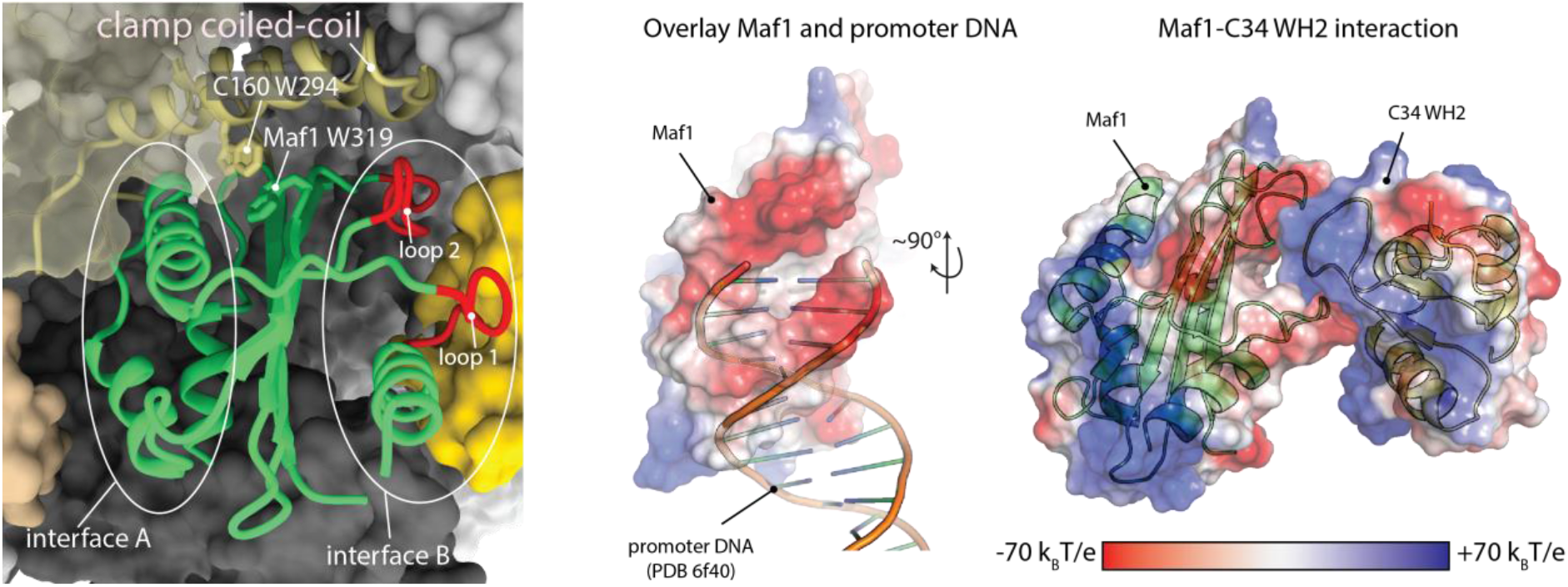
Details of the Maf1-Pol III interaction. Left: Close-up of the Maf1 binding site, with elements referred to in the text labelled. Right: Maf1 loops 1 and 2 mimic DNA and bind the mobile C34 WH2 domain.

### Conserved loops in Maf1 mimic DNA

Mapping the electrostatic potential onto the surface of Maf1 reveals a striking feature of interface B. Two loops, comprising residues 247-256 (loop 1) and residues 306-315 (loop 2), are negatively charged (Figure 2). Superposition of the Maf1-Pol III structure with the structure of the Pol III PIC in the open DNA state (PDB 6f40) further shows that the backbone of DNA partially overlaps with these loops, and that interface B is a near-perfect mimic of one turn of B-DNA backbone (Figure 2). The C34 WH2 binds this interface by inserting its positively charged ‘wing’ between the two loops. Comparison to *Homo sapiens* and *Citrus sinensis* Maf1 sequences and structures shows a good conservation of loop 1 and a conservation of the negative charge in loop 2. However, the *S. cerevisiae* sequence of loop 2 diverges from the consensus sequence of loop 2 compared to other species (Figure S3), which are even more negatively charged. Nonetheless, the negative charge of interface B is conserved, arguing that the mechanism of inhibition is also conserved (Figures S3 and S5).

### Maf1-Pol III-interface mutants impair transcription inhibition

To validate the structure, we generated mutants of Maf1 that we expected to show reduced binding to Pol III. The first mutant disrupts the aromatic stacking and hydrophobic interaction between the Pol III clamp coiled-coil and the Maf1 furrow (Maf1 L318S/W319S). The second mutation (Maf1 T33E/T34E) is expected to cause a clash between Maf1 and the Pol III protrusion. Lastly, we also prepared a previously described Maf1 D248A/D250A variant^18^ which reduces the negative charge in the DNA mimicking loop 1. We tested these proteins in an *in vitro* transcription assay as well as in pull-downs with recombinant C34^1-156^, which comprises the two N-terminal C34 WH domains. As expected, only the Maf1 D248A/D250A mutant showed reduced binding to C34^1-156^ (Figure 3). In our transcription assay, the D248A/D250A and L318S/W319S mutants show impaired transcription inhibition at increasing concentrations of Maf1, highlighting the importance of the aromatic stacking interaction and the DNA-mimicking acidic loops for efficient transcription inhibition. This is in line with the finding that the D248A/D250A mutant is completely inactive *in vivo*^18^. However, the T33E/T34E mutant inhibits transcription similarly as the wild-type protein, indicating that the bulkier amino acids at these positions can be accommodated.

**Figure 3.**
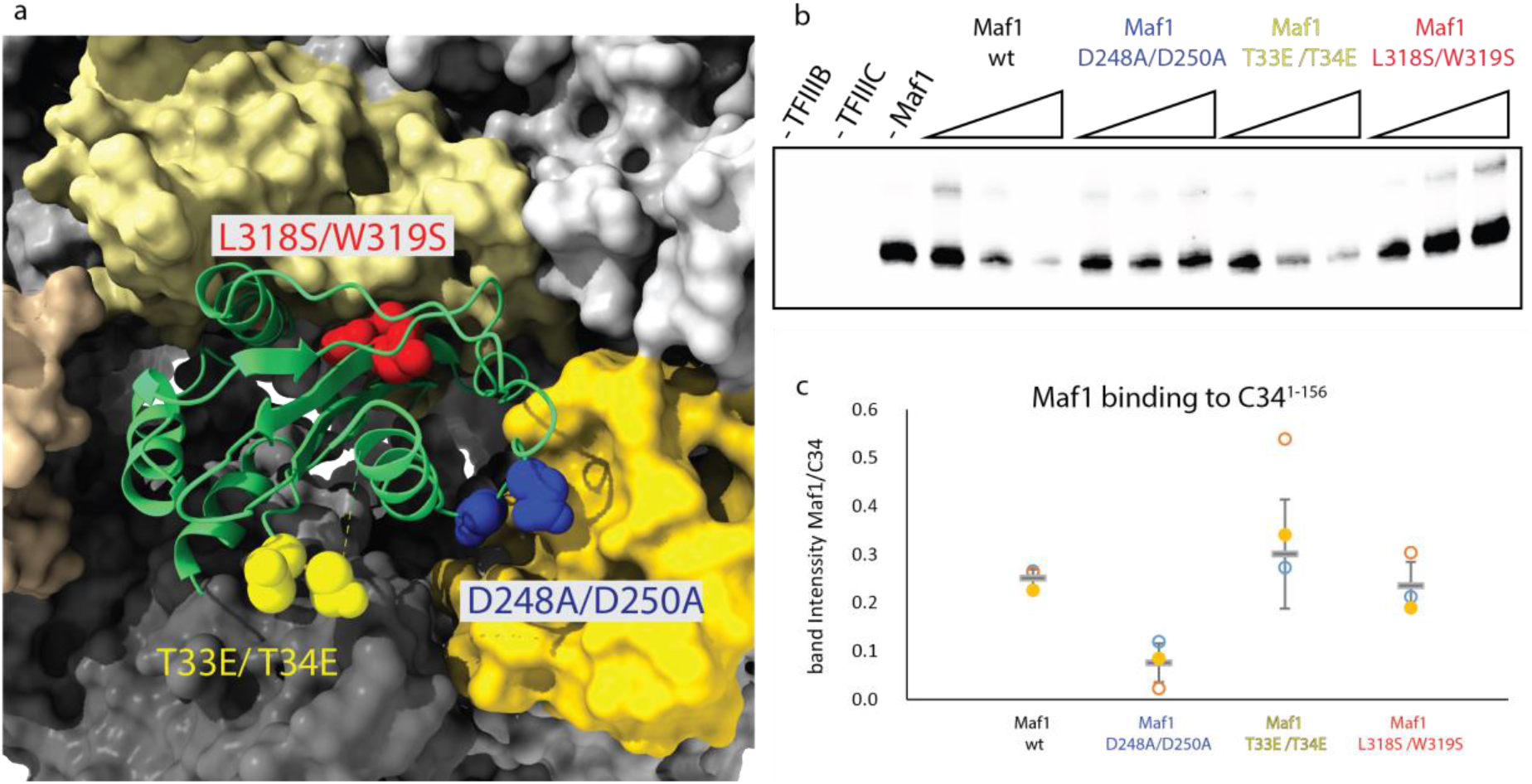
Maf1-Pol III-interface mutants impair transcription inhibition. a. Location of the introduced mutations. b: *In vitro* transcription assay comparing inhibition efficiency of Maf1-id wt and Maf1-id mutant proteins. c. Quantification of Maf1-id binding to C34 1-156 as determined from three independent pull-down. The centerlines represent the mean and vertical error bars represent the standard deviation of the mean.

To establish how binding of Maf1 triggers inhibition of transcription initiation, we overlaid the structure of Maf1-Pol III with the Pol III PIC (TFIIIB-Pol III) (Figure 4). The binding site of Maf1 overlaps extensively with the position of TFIIIB and promoter DNA in the PIC. More specifically, Maf1 clashes with the cyclin I domain of the Brf1 subunit of TFIIIB and with the double-stranded DNA near the upstream edge of the transcription bubble (Figure 4). This explains the finding that binding of Maf1 and TFIIIB are mutually exclusive^16^. Moreover, Maf1 induces allosteric changes that closely mimic the PIC state and sequesters residues that are involved in establishing contacts with promoter DNA. Specifically, the protrusion, the template strand pocket and the C34 WH2 ‘wing’ are all bound by Maf1, and the active site cleft is sealed off due to ordering of the C34 WH2 (Figures 1 and 2).

**Figure 4.**
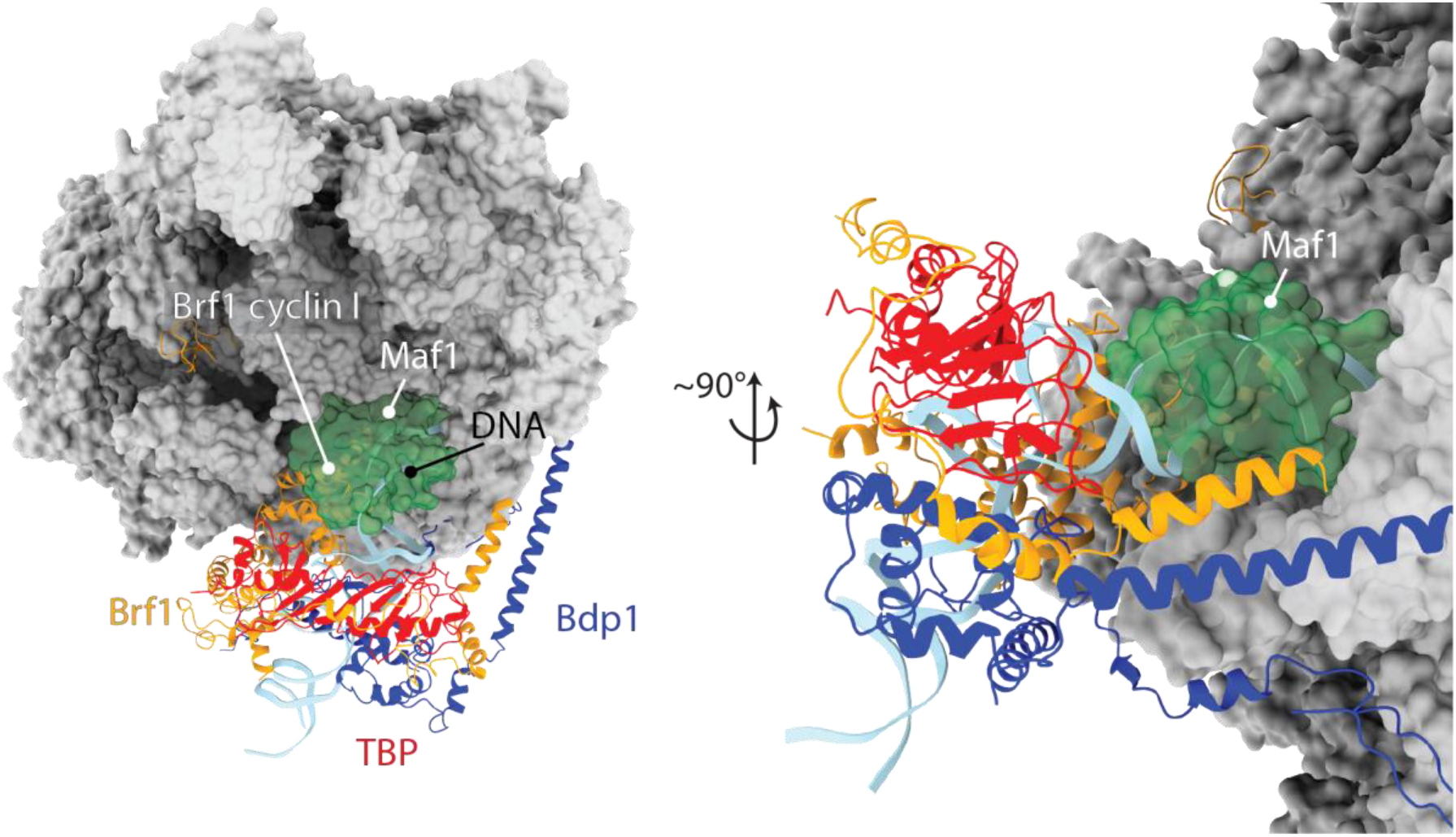
Maf1 clashes with TFIIIB and DNA in the PIC. Superposition of the Maf1-Pol III structure reported here and the Pol III pre-initiation complex (PDB 6f40) in two views. Maf1 is shown in green transparent surface and TFIIIB and promoter DNA in ribbon representation.

The C-terminus of Maf1 is acidic in different species but its sequence is poorly conserved^23^. This region of the protein is predicted to be disordered and was removed for X-ray structure determination of both *H. sapiens* and C. *sinensis* Maf1 proteins^16,17^. Interestingly, although the acidic C-terminus of *S. cerevisiae* Maf1 was retained in the protein used for Maf1-Pol III complex formation, it was not visible in the cryo-EM structure. The orientation of Maf1 relative to Pol III suggests that the C-terminus may project into the active site cleft (Figure S6). This might help to repel nucleic acids electrostatically and thereby prevent non-specific binding of Pol III to open chromatin. Whatever the function of the acidic C-terminus, the consequences of exchanging this domain with the corresponding region of *S. pombe* or *H. sapiens* Maf1 or of deleting the domain entirely are apparently subtle, since yeast cell growth on both fermentable and non-fermentable carbon sources and transcriptional repression appear normal in these mutants (Figure S7).

We also note that the regulatory internal region, that is disordered in our structure, and is subject to phosphorylation by TORC1, Sch9 and/or PKA^1,6^ is located on the solvent-exposed side of Maf1 (Figure S6). Maf1 can hence likely serve as a substrate to those kinases even in complex with Pol III, to allow rapid de-repression.

## DISCUSSION

The structure presented here differs from a previous low-resolution (~20 Å) cryo-EM study^16^, where Maf1 was reported to bind Pol III at a different location, namely on top of the clamp. The authors proposed that Maf1 achieves transcription repression by an allosteric mechanism, namely through rearrangement of the C82/C34/C31 heterotrimer complex, thereby locking Pol III in a conformation that is unable to bind TFIIIB. This is in contrast to the binding site and mechanism of inhibition demonstrated in this work. The near-atomic resolution of our cryo-EM map, and biochemical validation of our structure using structure-guided mutagenesis clearly establish that Maf1 binds between the clamp, wall and protrusion domains, and achieves transcription repression through direct competition with TFIIIB.

We note that in our Maf1-Pol III complex structure, the path of the upstream DNA is obstructed in a hypothetical Maf1-Pol III elongation complex. Yet in previous work Maf1 did not inhibit transcription from tailed templates or RNA-DNA-hybrids in factor-independent assays, and Maf1-Pol III-nucleic acid complexes could be purified in the absence of nucleotides^16^. We offer two explanations for these findings. First, the templates used in the study have short upstream DNA segments, and clashes between Maf1 and DNA are only expected with the upstream segment. Therefore, the template could be threaded into the active site, end-first, while Maf1 is still bound. Subsequent RNA synthesis could then displace Maf1 through the energy released during NTP hydrolysis in a similar fashion as TFIIIB is displaced during promoter escape. Second, it is possible that a second binding mode exists in which Maf1 does not obstruct the path of the DNA, perhaps mediated through disordered regions, as is the case for many protein-protein interactions governing the regulation of eukaryotic gene transcription^24^.

In summary, the results presented here provide a high-resolution structure of the Maf1-Pol III interaction, establish the exact mechanism of Pol III inhibition by Maf1, and rationalize previous biochemical data. Comparison of the *S. cerevisiae* Maf1 structure with available crystal structures from *H. sapiens* and *C. sinensis* suggest that this mechanism is conserved across kingdoms.

## Methods

### Protein purification

Pol III was purified as described^25^, but buffer exchanged into a buffer containing K_2_SO4 instead of (NH_4_)_2_SO4.

Maf1-id was expressed from a pETM11 vector in BL21 Star (DE3) pRARE *E. coli* cells in TB medium. Expression was induced with 500 μM IPTG at 25 °C overnight. Cells were pelleted for 5 min at 12,000 g and re-suspended in 3 mL lysis buffer (0.5M NaCl, 50 mM Tris pH 7.5, 2 mM β-mercaptoethanol (BME), 20% glycerol, 10 μg/mL DNase I, 1 x protease inhibitors (SIGMAFAST protease inhibitor cocktail EDTA free), 30 mM imidazole, 2 mM MgCl_2_) per gram of pellet. Cells were lysed in an Emulsiflex-C3 homogenizer and the lysate cleared by centrifugation for 1 h at 30,000 g. The supernatant was incubated with 5 mL Ni-NTA resin (Qiagen) for 2 h. Beads were recovered and washed with 100 mL His-A buffer (1 M NaCl, 50 mM Tris pH 7.5, 2 mM BME, 5% glycerol, 30 mM imidazole) and 50 mL His-A low salt (His-A but with 150 mM NaCl) and eluted with 50 mL His-B (200 mM NaCl, 2 mM BME, 5% glycerol, 300 mM imidazole). The eluate was supplemented with 1 mg of TEV protease and dialysed against 2 L of TEV-cleavage buffer (100 mM NaCl, 5 % glycerol, 1 mM DTT, 50 mM Tris pH 7.5) overnight. Cleaved protein was passed over 5 mL of Ni-NTA beads pre-equilibrated in TEV cleavage buffer. Full-length Maf1 and Maf1-id were further purified over a MonoQ HiTrap column and eluted with gradient from 100 mM NaCl to 500 mM NaCL over 20 CVs. Maf1-id was buffer exchanged in a spin concentrator to 100 mM NaCl, 1 mM TCEP, 15 mM HEPES, concentrated to 30 mg/mL and flash-frozen in liquid nitrogen. Maf1-id mutations were generated using the QuickChange mutagenesis protocol. Brf1-TBP and Bdp1 and TFIIIC were purified as described^21,26^.

C34^1-156^ was expressed from a pETM11 vector with an N-terminal his-tag and purified over Ni-NTA, as described for Maf1. The eluate was loaded onto a 5 mL Heparin FF column and eluted with a linear gradient from 100 mM NaCl to 1M NaCl over 20 CV. Peak fractions were concentrated and applied to a Superdex 75 gel filtration column pre-equilibrated in 150 mM NaCl, 20 mM HEPES pH 7.5 and 5 mM DTT. The purified protein was concentrated to 35 mg/mL and flash-frozen in liquid nitrogen.

### EM sample preparation

180 μg Pol III and a five-fold molar excess of Maf1-id were incubated overnight in binding buffer in a total volume of 48 μL(40 mM K_2_SO_4_, 15 mM HEPES pH 7.5 and 2.5 mM DTT). The sample was then diluted to 1.8 mL in the same buffer but containing 3 mM of BS3 crosslinker and 2.3 μM Maf1 to prevent dissociation. The sample was crosslinked for 2h at room temperature, quenched with 50 mM Tris pH 7.5, concentrated to ~60 μL and applied to a Superose 6 INCREASE 3.2/300 column equilibrated in EM buffer (150 mM (NH_4_)_2_SO_4_, 15 mM HEPES pH 7.5, 10 mM DTT). A peak fraction eluting at 1.4 mL was diluted to 0.15 mg/mL and used for grid preparation. 2.5 μL of sample were applied to Quanitfoil Cu 2/1 + 2 nm Carbon grids that were glow-discharged in a Pelco EasyGlow instrument. Access sample was blotted away in a Vitrobot Mark IV set to 4 °C and 100 % humidity for 6 seconds with a blot force of 2 and the grid was plunge-frozen in liquid ethane.

### Electron microscopy

Cryo-EM data was collected on a Titan Krios microscope with a Gatan Quantum energy filter and a K2 Summit direct detector in counting mode at a magnification of 130 000x and a pixel size of 1.041 Å /px. 10520 movies with 40 frames each and an accumulated dose of 60.5 e^-^/Å^2^ were collected, acquiring 8 images per hole.

Movies were pre-processed on the fly using Warp 1.05^27^. The model parameters for motion correction and CTF estimation were set to 5×5×40 and 5×5×1, respectively. Particles were picked with BoxNet2_20180602 without re-training, using an expected particle diameter of 250 Å. Particles were extracted in a 360 pixel box, normalized and the contrast inverted. Filters in Warp were set to exclude particles from micrographs with lower than 6 Å estimated resolution based on CTF fitting, with a larger than 1.7 Å movement per frame in the first third of the movie, and with fewer than 50 particles per micrograph. However, very few micrographs did not satisfy these criteria (125 excluded micrographs, ~1% of the collected data), indicating high quality of the sample. Warp was also used for denoising of micrographs as shown in Figure S1.

### Particle classification

Due to the large number of particles (1 688 795), the dataset was divided into four batches and each was classified by 3D classification in RELION 3.0 using a 60 Å low-pass filtered model of apo Pol III (PDB 5fj9) as the reference. The best class of each batch was retained, yielding a dataset of 728 000 particles. All batches were combined and refined.

The resulting map showed density for Maf1 and an adjacent density corresponding to C34 WH2, but at a lower threshold. Using a classification focused on this region the Maf1-C34WH2 density could be improved (305 000 particles). Finally, a masked classification focusing on the stalk, trimer and clamp separated an open clamp state and a closed clamp state. Density for Maf1 was poor in the open clamp state, but clear separation of beta-sheets and sidechain density were visible in the closed clamp state.

Two rounds of CTF refinement improved the resolution from 3.4 Å to 3.25 Å. Starting from the 305 000 particle set, we also performed MultiBody refinement implemented in RELION 3, using two masks. The first covered the core of Pol III and the heterodimer, and the second covered the stalk, clamp, heterotrimer and Maf1. The resulting bodies were sharpened using the relion_postprocess program, yielding maps at 3.34 Å (core) and 3.74 Å (stalk-trimer-clamp) resolution. This significantly improved side-chain density in the heterotrimer and clamp.

### Model building and refinement

For Model building, PDB 6eu3 (apo-Pol III closed conformation at 3.3 Å resolution) was used as a starting point and combined with the C34 WH2 domain from PDB 6f40. A homology model of yeast Maf1 was generated with PHYRE2^28^ and fitted into the density. The model was manually adjusted in COOT^29^.

To improve model coordinates, we used the maps obtained by MultiBody refinement and corrected minor sequence register shifts that were present in the starting model and could be detected due to the higher quality of the MultiBody maps. Progress of model building was monitored by real-space refinement in PHENIX^30^ of partial models, namely the heterodimer, the heterotrimer-clamp region, and Maf1 against respective maps obtained from MultiBody refinement. Finally, the refined models were combined and refined in another round of real-space refinement against the locally filtered 3.25 Å consensus map obtained using the relion_postprocess program. MolProbity and PHENIX model validation statistics show good stereochemistry and excellent correlation between the map and the model (correlation coefficient of 0.81, and a FSC0.5 of the simulated model map with the experimental map of 3.1 Å, see Supplementary Figure 1).

### Maf1-C34 pulldown

His-tagged C34^1-156^ (180 μg) was incubated with an equimolar amount of wild-type and mutant Maf1 proteins for 1h, and then diluted tenfold in IP buffer (50 mM NaCl, 20 mM HEPES pH 7.5 and 5 mM DTT). The sample was applied to a small gravity column containing 50 μL of Ni-NTA beads which was then washed with 1.5 mL of IP buffer. Proteins were eluted in 100 μL of elution buffer (300 mM imidazole, 200 mM NaCl, 50 mM Tris pH 7.5) and analysed by SDS-PAGE. Binding was quantified by measuring band intensities using the ImageLab software, and the relative intensity of Maf1 bands to C34 bands in the eluted fraction was calculated. The experiment was repeated three times using the same protein batches.The mean values and standard deviations were calculated in Microsoft Excel.

### *In Vitro* Transcription

15-μl transcription reactions contained 2 pmol tRNA^His^ hybridized DNA oligonucleotides (template-strand 5’AAAATGCCATCTCCTAGAATCGAACCAGGGTTTCATCGCCACA-CGATGTGTACTAACCACTATACTAAGATGGCGACTTTTCAATGGAGAACTGTTGTATTACGGGCTCGAGTAATAC 3’ and non-template strand 5’-GTATTACTCGAGCC-CGTAATACAACAGTTCTCCATTGAAAAGTCGCCATCTTAGTATAGTGGTTAGTACACATCGTTGTGGCCGATGAAACCCTGGTTCGATTCTAGGAGATGGCATTTT-3’), 4 pmol TFIIIC, 4 pmol TFIIIB, 3 pmol Pol III, and either 8, 16 and 24 pmol Maf1-id wt or mutants in 20 mM HEPES pH 7.5, 60 mM (NH_4_)_2_SO_4_, 10 mM MgSO_4_, 10% glycerol, 10 mM DTT, 120 μM of each ATP, GTP, and CTP as well as 10 μCi of [α-^32^P]UTP (Hartmann). Maf1 wt or mutant proteins were added to the DNA before TFIIIC as well as pre-incubated with Pol III. *In vitro* transcription was performed at RT for 1 hr before phenol/chloroform extraction and precipitation. The RNA was then subjected to electrophoresis in denaturing 12% polyacrylamide gels. The gels were exposed to phosphorimaging screens (Fujifilm) for capturing digital images.

### Yeast Phenotypes and Northern blotting

Standard cloning methods were used to delete the non-conserved acidic C-terminus of full-length *S. cerevisiae* Maf1 in pRS314 (Moir et al., 2006) and to substitute the corresponding regions of *S. pombe* and human Maf1. After transformation into a W303 *maf1*Δ::*natMX* strain, growth was monitored on media containing glucose or glycerol and northern analysis was performed as reported previously (Moir et al., 2006).

## DATA AVAILABILITY

The cryo-EM map of the Pol III-Maf1 complex has been deposited to the Electron Microscopy Data Bank (EMDB) under the accession code EMD-XXXX. The coordinates of the corresponding model has been deposited to the Protein Data Bank (PDB) under accession code XXXX

## ACKNOWLEDGEMENTS

This work was supported by an ERC Advanced Grant (ERC-2013-AdG340964-POL1PIC) to C.W.M. and R.W., a National Institutes of Health Grant (GM120358) to I.M.W. and an EMBL International PhD program award to M.K.V. We thank M. Girbig for help with transcription assays and critical reading of the manuscript and F. Weis for EM support. We are thankful to T. Hoffmann and J. Pecar for maintaining the high performance computing environment for EM data processing at EMBL.

## AUTHOR CONTRIBUTIONS

The project was conceived by C.W.M., I.M.W and R.D.M and supervised by C.W.M.. M.K.V. designed, carried out and analyzed experiments, data processing and model building. F.B performed transcription assays. W.J.H.H. collected cryo-EM data. R.W. was responsible for yeast fermentation. M.K.V. and C.W.M. prepared the manuscript with input from the other authors. R.D.M. performed functional analysis of the acidic tail.

## COMPETING INTERESTS

The authors declare no competing interests.

## Supplementary Figures

**Figure S1.**
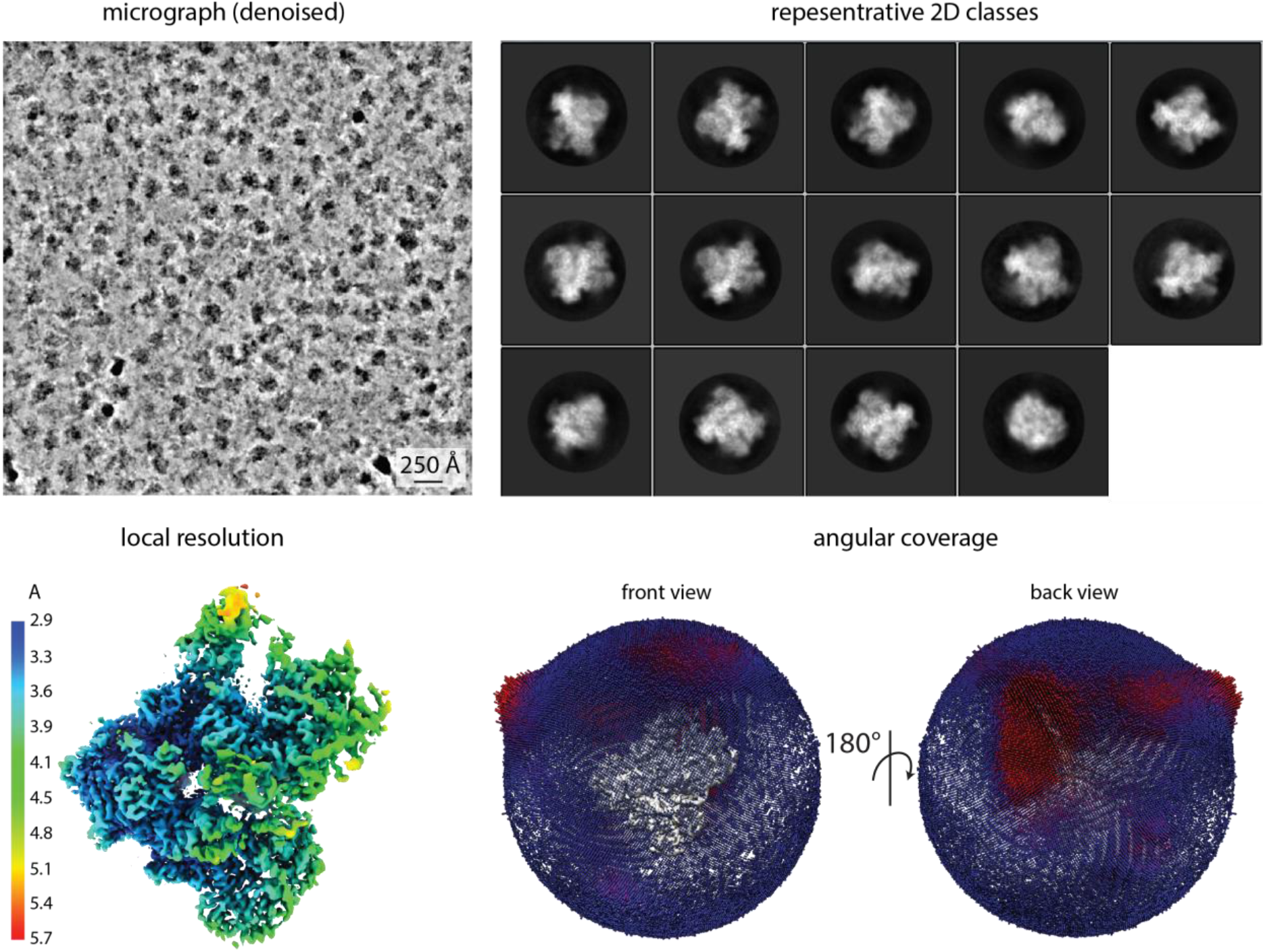
Cryo-EM data quality. Representative micrograph denoised using Warp^27^, 2D classes, EM density colored by local resolution, and angular distribution.

**Figure S2.**
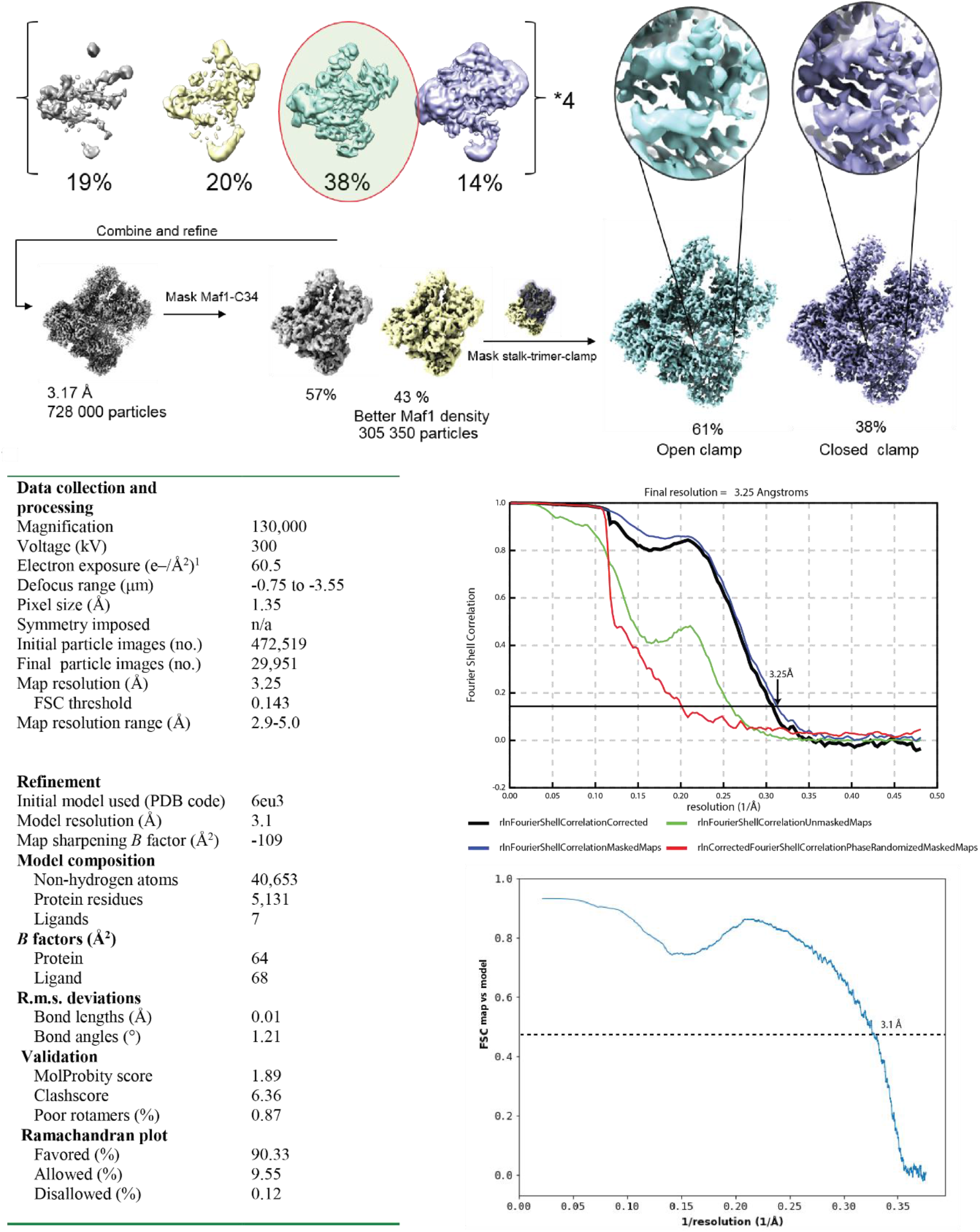
Particle classification, data collection and refinement statistics. Top panel: Micrographs were divided into four batches and classified using 3D classification in RELION with a 60 Å low-pass filtered map of Pol III as a reference. The best class of each batch was retained, batches were combined and refined. Focused classification using a mask on Maf1 yielded one class with improved Maf1 occupancy. Lastly, focused classification using a mask on the stalk-heterotrimer-clamp module separated an open clamp state with poorly resolved Maf1 density from a closed-clamp state with clear separation of the Maf1 β-strands and side-chain density. Bottom panel: Data collection and refinement statistics. Plots show the FSC curve of the two independent half-maps (top) and the FSC curve of our experimental density vs density simulated from the model.

**Figure S3.**
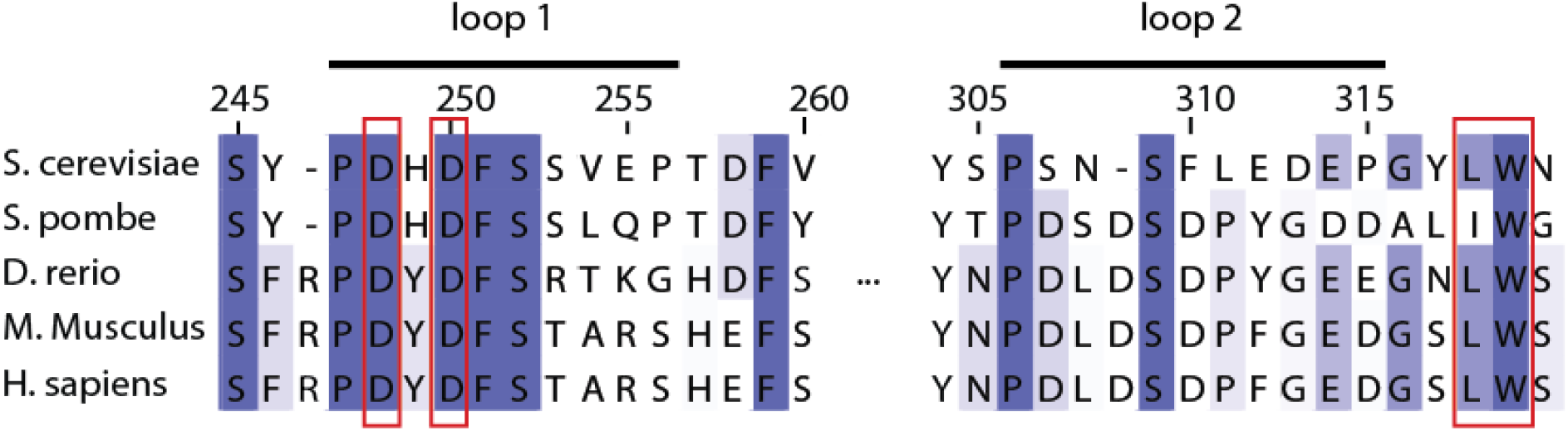
Multiple sequence alignment of Maf1, showing sequences of loop 1, loop 2 and W319 from distantly related species. Functionally important sites based on mutant studies are boxed in red.

**Figure S4.**
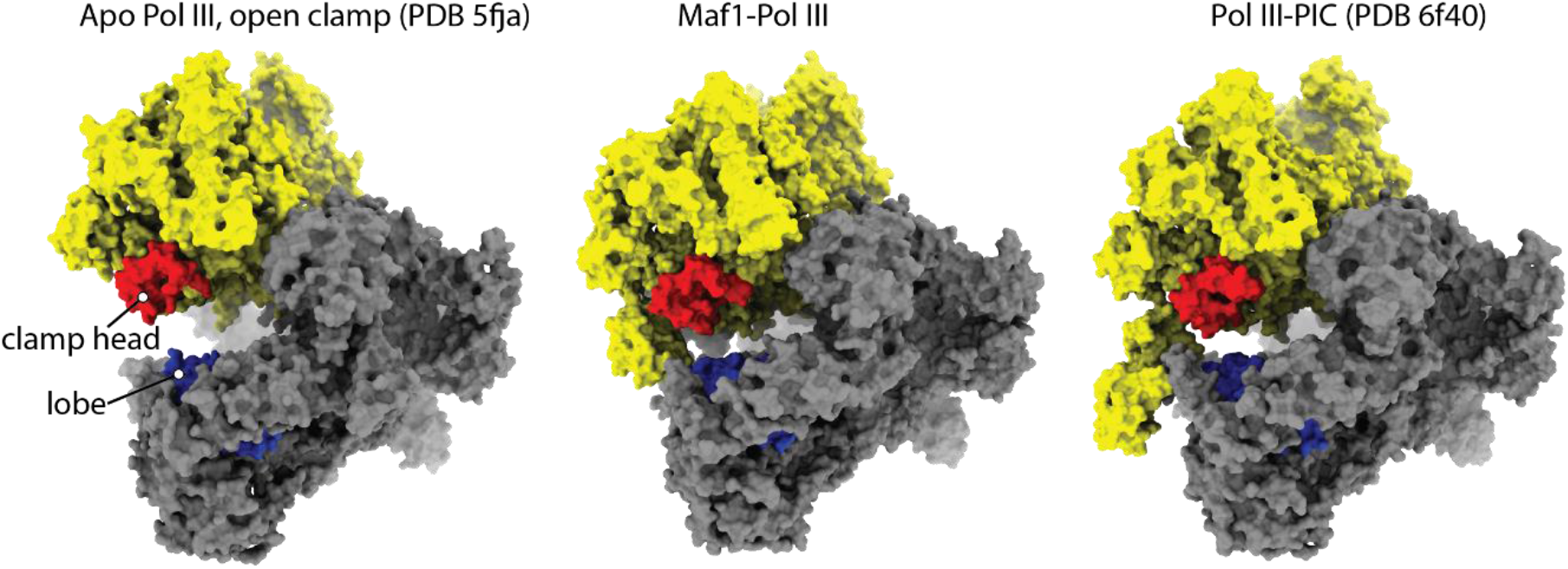
Comparison of the Pol III conformation in apo-Pol III, Maf1-Pol III and the Pol III-PIC. Maf1-Pol III adopts a similar conformation as the Pol III-PIC (note the distance between the lobe (blue) and clamp head (red)).

**Figure S5.**
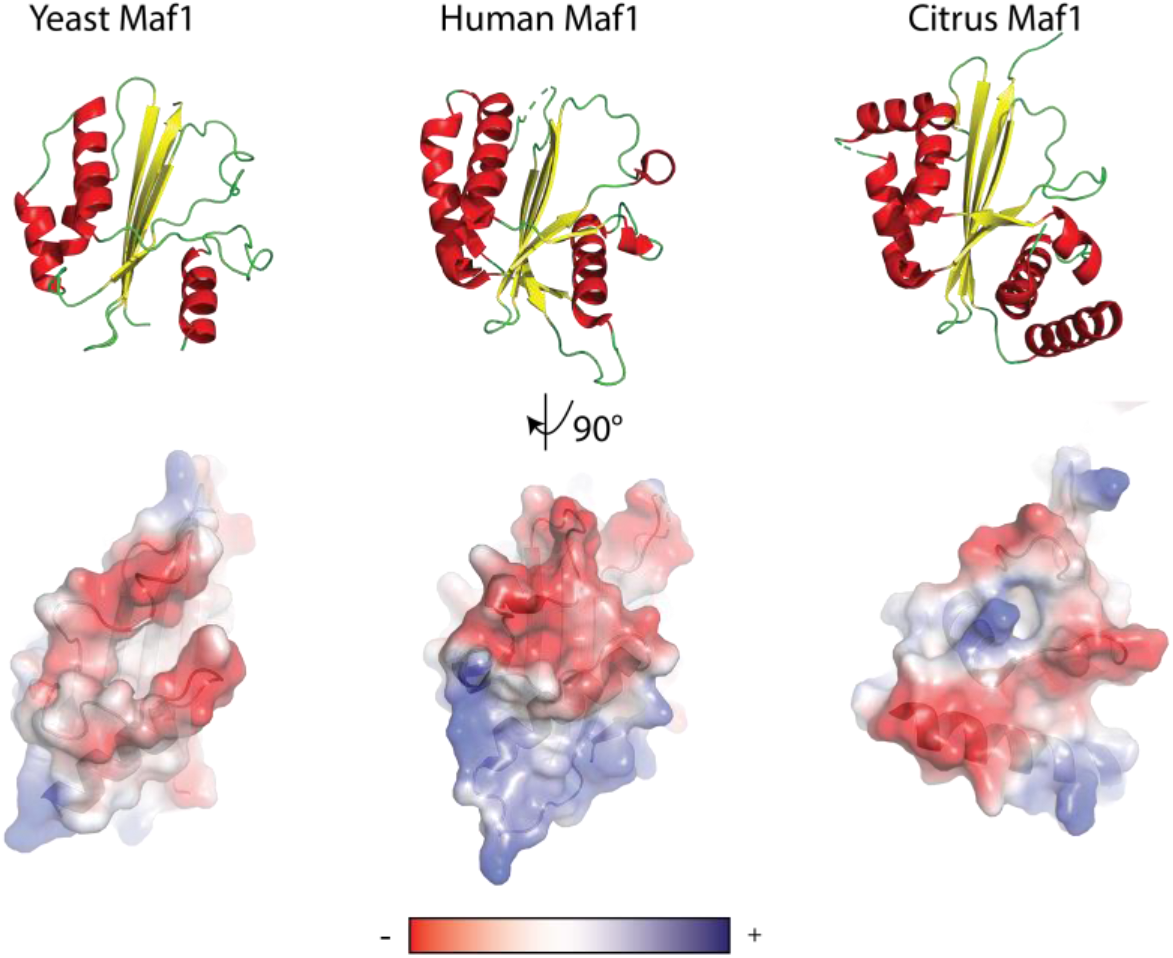
Comparison of Maf1 structures from *S. cerevisiae, H. sapiens* and *C. sinensis*. Top panel: Structures shown in ribbon representation with α-helices colored in red and β-sheets colored in yellow. Bottom panel: surface representation colored by electrostatic potential from negative (red) to positive (blue) potential.

**Figure S6.**
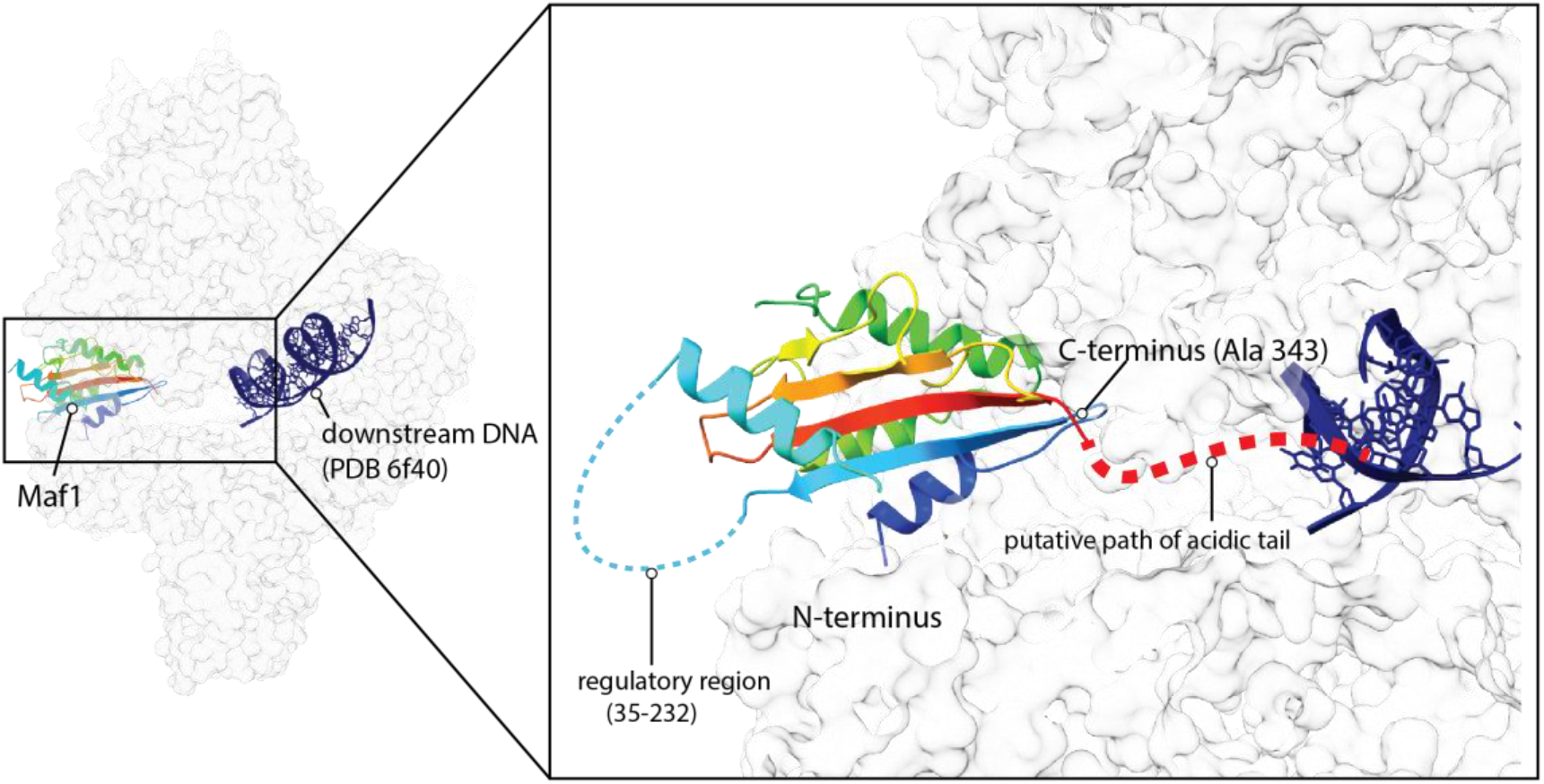
A model of the disordered regions in Maf1. The phospho-regulatory region of Maf1 is accessible in the Maf1-Pol III structure, whereas the acidic tail emerges in the direction of the DNA binding cleft. The downstream DNA (PDB 6f40) has been superimposed with the Maf1-Pol III structure to help visualization. Maf1 is colored from N-terminus (blue) to C-terminus (red). Putative paths of the disordered regions are indicated, with the internal, phospho-regulated region located away from the Pol III interface, whereas the acidic C-terminal tail projects towards the DNA binding cleft, where it might help to repel nucleic acids.

**Figure S7.**
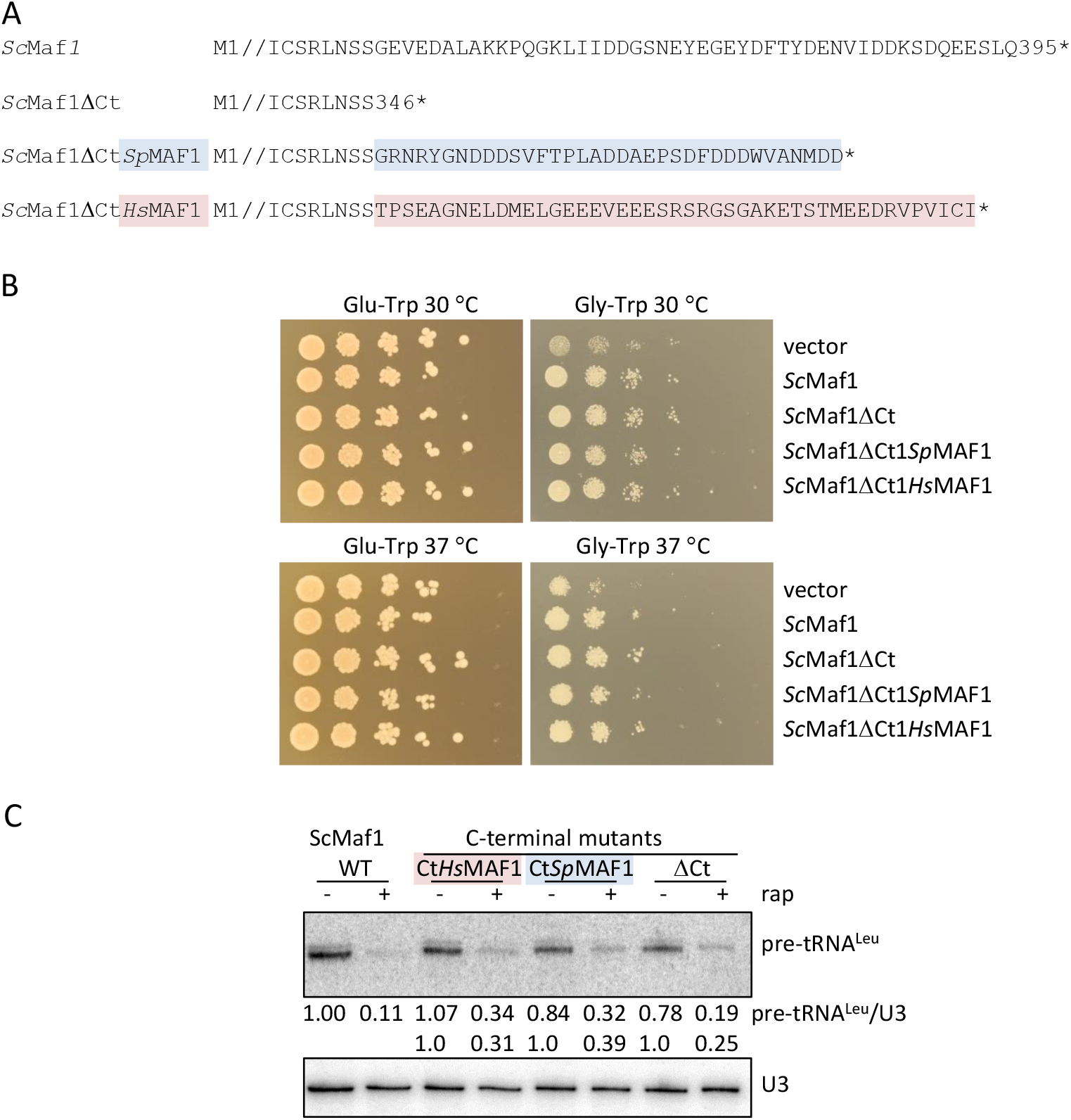
The C-terminus of ScMaf1 is not obligatory for Maf1 function. (A) Amino acid sequences of wild-type ScMaf1 and the various ScMaf1 C-terminal mutants are shown from the end of conserved domain C (Pluta et al., 2001) through the acidic terminal region (where present). Amino acid sequences from residue 2 through 326 are represented by //. ScMaf1ΔCt terminates at residue 346 while ScMaf1ΔCtSpMAF1 and ScMaf1 ΔCtHsMAF1 proteins contain the terminal amino acids from *S. pombe* (35 residues, colored in blue) and human (45 residues, colored in pink) MAF1 proteins appended to ScMaf1ΔCt. (B) The respiratory defect of the *maf1Δ::natMX* vector only strain, poor growth on glycerol, is rescued by wild-type ScMaf1 and the three ScMaf1 C-terminal variants. Ten-fold serial dilutions of *maf1Δ:: natMX* strains containing pRS314 vector, pRS314ScMaf1 or pRS314ScMaf1 C-terminal variants grown at 30 and 37°C on media with glucose (left panels) and glycerol (right panels) as the carbon source. (C) Northern analysis of Pol III transcription and repression shows that the C-terminus of ScMaf1 is not required for the Maf1 repression function in yeast. *maf1Δ::natMX* strains containing pRS314ScMaf1 or pRS314ScMaf1 C-terminal variants (ScMaf1ΔCt, ScMaf1ΔCtSpMAF1 and ScMaf1 ΔCtHsMAF) were treated with rapamycin or drug vehicle for 1 h. The relative level of Pol III transcription is reported by the amount of pre-tRNA^Leu^ transcript normalized to U3 snRNA, expressed relative to the untreated wild-type strain and indicated below each lane.

## Notes

#### Summary of Updates

Formatting of accented characters has been fixed

